# *MS.liverK:* an R package for transcriptome-based computation of molecular subtypes and functional signatures in liver cancer

**DOI:** 10.1101/540005

**Authors:** Florent Petitprez, Léa Meunier, Eric Letouzé, Yujin Hoshida, Augusto Villanueva, Josep Llovet, Snorri Thorgeirsson, Xin Wei Wang, Wolf H Fridman, Jessica Zucman-Rossi, Aurélien de Reyniès

## Abstract

**Summary:** Liver cancer is a highly heterogeneous disease in terms of etiology, tissue and cellular morphology, tumor molecular characteristics, microenvironment composition and prognosis. Several studies, based on tumor gene-expression profiling (GEP) data, have dissected the molecular heterogeneity of liver cancer. They resulted in various tools, either delineating homogeneous tumor subtypes or calculating molecular scores of prognostic or biological functions. Here, we present MS.liverK, an easy-to-use R package providing a comprehensive implementation of these tools, for research use.

**Availability and implementation:** The *MS.liverK* R package is available from GitHub (https://github.com/cit-bioinfo/MS.liverK).

## 1. Introduction

Primary liver cancer is the third most deadly cancer worldwide, with Hepatocellular carcinomas (HCC) accounting for 85%-90% of the cases. The molecular heterogeneity of liver cancers has motivated numerous transcriptome-based studies, yielding gene-signatures and algorithms for molecular subtyping, prognostic prediction and functional scores computation. The use of these tools is of great interest for researchers dedicated to liver cancer study, but is very limited due to their dispersion.

Here, we introduce MS.liverK (Molecular Signatures in Liver Cancer), an R package reimplementing these tools. MS.liverK takes as input a (log2 scale) transcriptome matrix of liver cancer samples, either microarray-or RNA-Seq-derived. It also includes graphical functions, allowing to easily visualize the outputs.

## 2. MS.liverK

After a careful review of the literature, we identified 5 GEP-based molecular subtyping systems of HCCs published in the last 15 years (Lee *et al.*, 2004; Boyault *et al.*, 2007; Yamashita *et al.*, 2008; Chiang *et al.*, 2008; Hoshida *et al.*, 2009). Lee classification (Lee *et al.*, 2004) was established on a belgo-chinese cohort of 91 HCC samples; it is made of 2 classes (A/B), related to proliferation and survival. Lee subtypes have been further refined in 4 groups using Alpha-fetoprotein (AFP) marker, improving association to survival. Boyault classification (Boyault *et al.*, 2007) was established on a French cohort of 57 HCCs including patients from various geographic origins; it is made of 6 classes (G1 to G6) related to TP53 and CTNNB1 mutations, proliferation, HBV infection and prognosis. Yamashita classification (Yamashita *et al.*, 2008) is based on two markers, EPCAM and AFP, and defines four groups related to differentiation and prognosis. Chiang classification (Chiang *et al.*, 2008) was built using 91 HCV infected HCCs; it identifies 5 classes related to CTNNB1 mutation, proliferation, inflammation and polysomy of the chromosome 7. Hoshida classification (Hoshida *et al.*, 2009) is based on the meta-analysis of 9 cohorts totaling 603 HCC samples; it contains 3 classes (S1/S2/S3) related to proliferation, differentiation and survival. MS.liverK implements all of the above classification systems from transcriptomic data; it also includes a conversion function to allow using different GEP platforms (Fig. S1A).

To ensure classifiers were adequately assigning classes to samples, we ran them on the data from the samples used by the four teams to define their subgroups. We then compared MS.liverK classes to the ones they originally established and found a very good correspondence (Fig. S1B): Chi-squared test p-values for Lee, Boyault, Chiang and Hoshida and Roessler classifications were all < 2.2 e-16, with accuracy rate of respectively 94.5%, 98.2%, 100%, 93.8% and 90.0%.

MS.liverK implements the prognostic prediction algorithm published by (Nault *et al.*, 2013), which is so far the most extensively validated prognostic score for hepatocellular carcinomas (HCC).

Several teams have published gene-signatures related to biological functions/pathways involved in liver cancer oncogenesis. These gene-signatures relate to TGFB1 signalling (Coulouarn *et al.*, 2008), MET pathway (Kaposi-Novak *et al.*, 2006), stemness (Oishi *et al.*, 2012; Yamashita *et al.*, 2008), EPCAM (Yamashita *et al.*, 2008) and hypoxia (van Malenstein *et al.*, 2010). MS.liverK computes all the corresponding scores.

We applied MS.liverK to the TCGA LIHC RNA-seq dataset (n=286). The graphical output shows a striking concordance across the various subtyping systems, the prognostic score and the functional scores (Figure 1). This observation is expected, given that molecular subtypes should represent homogeneous types of tumors with specific biological and prognostic characteristics.

**Fig. 1.**
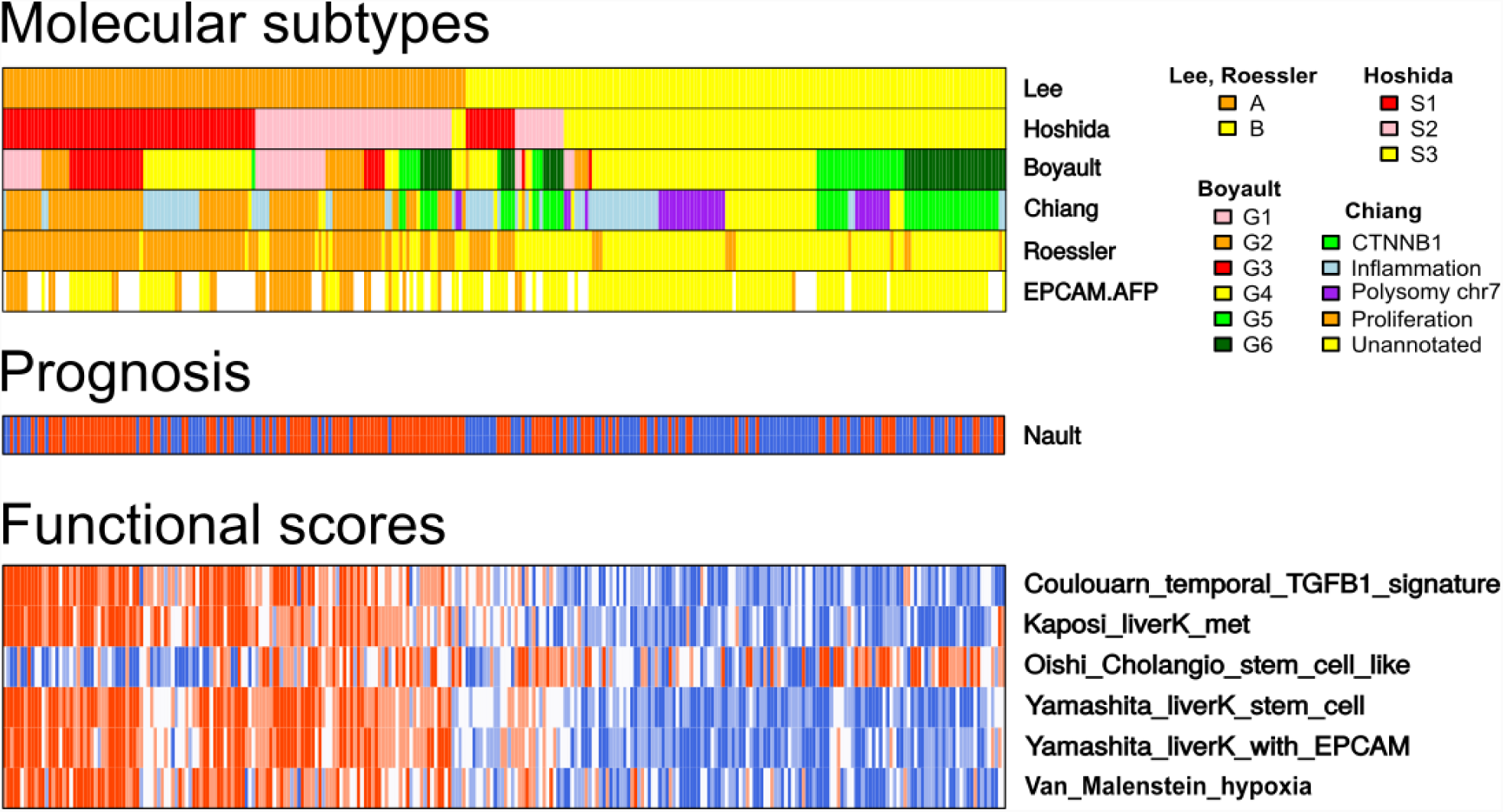
MS.liverK graphical output. Graphical output of MS.liverK illustrated on the TCGA LIHC transcriptome dataset. Columns represent samples, rows represent molecular subtypes (upper panel), prognostic scores (middle panel) and functional scores (lower panel).

## 3. Software implementation and use

The package has been entirely written using R language. It includes pre-packaged data from Gene Expression Omnibus (GEO), accession number GSE20238 (Mínguez *et al.*, 2011), to be used as example data. A vignette describing the use of the package is available on GitHub (https://github.com/FPetitprez/MS.liverK/blob/master/vignettes/vignette.pdf). On a standard computer, the computation of all subtypes and scores takes a couple of minutes.

