## Supplementary material for "*MS.liverK:* an R package for transcriptome-based computation of molecular subtypes and functional signatures in liver cancer": Figure S1

### A - Analytical workflow

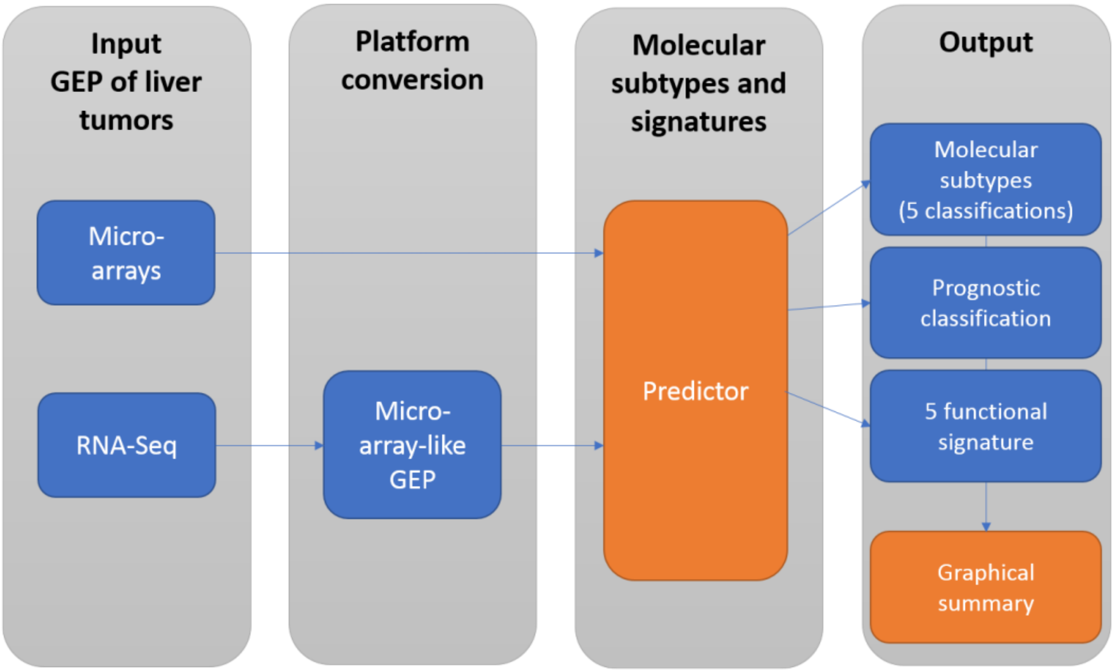


### B - Validation on original series

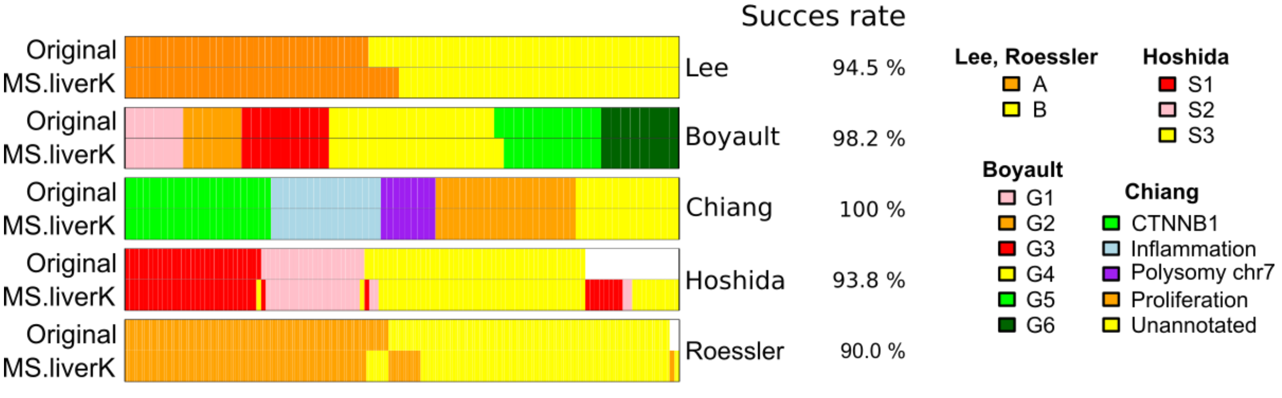


Fig. S1: A. Analytical workflow: (i) MS.liverK takes liver tumors GEP as input -several GEP platforms are supported-, (ii) when needed GEP are converted to microarray-like GEP, (iii) Molecular subtypes, prognostic and functional scores are calculated (iv) textual and graphical outputs are provided. B. Application of MS.liverK to TCGA LIHC cohort and performance of MS.liverK molecular subtyping features with respect to the original data of each classification system.
